# Single injection of very mild dose BOTOX in the vastus lateralis improves testicular spermatogenesis and sperm motility in ageing experimental mice

**DOI:** 10.1101/2021.10.24.465594

**Authors:** Risna Kanjirassery Radhakrishnan, Sowbarnika Ravichandran, Aishwarya Sukesh, Balamuthu Kadalmani, Mahesh Kandasamy

**Author notes:** Address for correspondence Dr. Mahesh Kandasamy, PhD., UGC-Assistant Professor, Department of Animal Science, School of Life Sciences, Bharathidasan University, Tiruchirappalli – 620024, Tamilnadu, India.

## Abstract

Acetylcholine (ACh), a key neurochemical messenger that plays key roles in neuroplasticity and muscle contraction. While ACh is important for the physiological function of the testis, abnormal levels of ACh cause testicular atrophy and male infertility. BOTOX is a therapeutic form of the botulinum neurotoxin that blocks the excessive release of ACh at the neuromuscular junction. Previously, repeated intracremasteric injections and slight overdose of BOTOX have been reported to induce adverse effects in the testicular parameter of experimental rodents. However, a mild dose of BOTOX is highly beneficial against skin ageing, neurological deficits, overactive urinary bladder problems, testicular pain and erectile dysfunctions. Considering the facts, the possible therapeutic benefit of BOTOX on the testis might be achieved via its minimal dose and indirect mode of action rather than repeated high quantity in the local supply. Therefore, we revisited the effect of BOTOX but with a trace amount injected into the vastus lateralis of the thigh muscle, and analyzed histological parameters of testis and quality of semen in ageing experimental mice. Experimental animals injected with 1 U/kg bodyweight of BOTOX showed enhanced spermatogenesis in associations with increased activities of key antioxidants in the testis, leading to increased total sperm count and motility. This study signifies that a mild intramuscular dose of BOTOX can be considered as a potential treatment strategy to manage and prevent male infertility.

## Introduction

Acetylcholine (ACh) is a key neuromodulator of the cholinergic system and biochemical regulator of muscle contraction [1]. ACh plays an important role in steroidogenesis and spermatogenesis in the testis [2,3]. Abnormal ageing, psychological complications, neuropathogenic and abnormal metabolic conditions associated with elevated levels of ACh and altered cholinergic signaling have been reported to induce erectile failure and testicular defects leading to endocrine imbalance and infertility [4–7]. Therefore, blockade of excessive ACh release can be proposed to restore male reproductive physiology that is lost upon ageing and various diseases. Botulinum toxins are a class of fatal proteins, mainly produced by *Clostridium botulinum*, cause muscle paralysis as they act by binding presynaptically to high-affinity recognition sites on the cholinergic nerve terminals and effectively prevents the exocytosis of ACh vesicles at the neuromuscular junction [8–10]. Notably, a trace amount of purified botulinum toxin has been proposed to yield long lasting anti-ageing therapeutic benefits and mitigate neuromuscular defects in various illnesses [8,9,11]. However, few toxicological reports indicated that cremasteric injection of botulinum toxin (10, to 40 U/kg) induces cell death and impairs spermiogenesis in the testis of experimental animals [12,13], while a recent report indicates that intrascrotal injection of botulinum toxin in adult male did not induce any adverse effect [14]. Nevertheless, ample scientific evidence unequivocally points towards the positive episodes of a very mild dose of therapeutic botulinum toxin such as BOTOX® against various clinical complications including movement disorders, cognitive deficit and behavioral abnormalities [8,9,11,15–17]. Eventually, therapeutic forms of botulinum toxins have been used as a potent treatment agent against skin ageing, hyperhidrosis, chronic migraine, strabismus, movement disorders, overactive urinary bladder problems, testicular pain and erectile dysfunctions [8,9,18–20]. Notably, mild doses of therapeutic botulinum toxins have been reported to mitigate oxidative stress and facilitate cytoprotection in various tissues including the brain [11,21–23]. Considering the facts, we speculate that the reported adverse effect of botulinum toxin on the testicular parameters might be largely due to a slightly high dose at the proximal site and repeated injections. Thus, we revisited the effect of BOTOX^®^ on the histological parameter of testis and quality of semen in ageing experimental mice. In this study, ageing experimental male mice were injected with the single dose of 1 U BOTOX^®^ per Kg bodyweight into the vastus lateralis of the thigh of experimental mice. After a four weeks’ time interval, experimental mice were sacrificed and histological parameters of spermatogenesis, sperm count and motility were performed in corroboration with biochemical assessments of antioxidant levels in testis.

## Materials and Methods

### Injection of BOTOX^®^ in experimental animals

For this study, 7-8-months-old (N=12) male BALB/c mice were randomly divided into two groups, namely control group (N□=□6) and BOTOX® treated group (N□=□6). BOTOX^®^ (Allergan, Dublin, Ireland) was dissolved in sterile saline. Experimental mice in the test group have received a single intramuscular injection of BOTOX^®^ at 1U per kilogram (Kg) bodyweight (BW) in the vastus lateralis muscle of the thigh as previously described [11,23]. An equal volume of sterile saline was injected to each mouse in the control group. Four weeks later, each mouse was sacrificed, the left testis was dissected out and processed for histological examination of spermatogenesis, while the right testis and cauda epididymis were processed for biochemical analysis of oxidative stress and antioxidant levels, and sperm analysis respectively. All the experiments were performed in accordance with the approval of the institutional animal ethical committee (IAEC) under the regulation of the Committee for the Purpose of Control and Supervision of Experimental Animals (CPCSEA), at Bharathidasan University (Reference No: BDU/IAEC/P272018, Date: 07.08.2018).

### Sperm count and motility

To assess the sperm parameters, the cauda epididymis was removed from each animal and minced in 1% bovine serum albumin (BSA) prepared with 1 X phosphate buffered saline (PBS) at 37ºC. The resulting epididymal fluid was diluted in the same solution (1:4), placed in the Neubauer chamber. The number of sperms and their degree of motility were estimated using an inverted phase contrast microscope (DMi1, Leica Microsystems, Germany). The total sperm count was calculated using the formula as Sperm count/ml = Mean sperm count x dilution factor x 10^4^. The motility of the sperms was graded as total and progressive motility, and percentage motility was estimated as previously described [24]. Next, smears of the epididymal fluid were prepared on the microscopic slides (Borosil, India) and stained using a solution containing haematoxylin (H) and eosin (E) (Nice Chemicals, India). The H and E-stained sperms on the slides were mounted using Dibutylphthalate Polystyrene Xylene (DPX) (Merck, Germany) and dried overnight at room temperature. The morphology and structure of the sperms were evaluated under the light microscope equipped with a camera and the software programme Leica Application Suite (DM750, Leica Microsystems, Germany).

### Histological evaluation of seminiferous tubules and spermatogenesis

The testes were fixed with 10% neutral buffered formalin (NBF), dehydrated using varying concentrations of ethanol (70%, 80%, 90%, 100%) followed by clearing using xylene, after which the tissues were infiltrated and embedded in paraffin wax. The wax embedded tissues were cut into cross-sections of 10 μm thickness using a rotary microtome (Weswox, India). The testicular sections were placed in silane coated microscopic glass slides and stained with H and E (Nice Chemicals, India). The histological sections were examined and photodocumented using a light microscope. The total number of the cross section of seminiferous tubules were counted in 10 non-serial cross sections of the testis in each animal from the control and treatment group. The diameters of the cross section of seminiferous tubules and lumens were measured as previously described [25]. Further, the total number of primary spermatocytes, secondary spermatocytes, round spermatids, and elongated spermatids were systematically quantified in seminiferous tubules per animal. The data on the estimated number of cells are represented per seminiferous tubule [26].

### Estimation of enzymatic activities of testicular antioxidants

The testicular tissue samples were homogenized with radioimmunoprecipitation assay (RIPA) buffer (Thermo scientific) and centrifuged at 12000 rpm for 20 minutes at 4ºC. The supernatant was collected and protein estimation was done by the method described by Lowry et al [27]. The total protein isolates were subjected for the measurement of key antioxidant enzymes as previously described [11]. In order to measure the activity of superoxide dismutase (SOD), the tissue homogenates were mixed with ice-cold ethanol and chloroform and centrifuged for 15 minutes at 12,000 rpm and the supernatants were collected. Further, 0.1 M tris buffer (pH 8.2) was added to the supernatant, and the reaction was initiated with the addition of 2.64 mM pyrogallol. As one unit SOD activity represents the amount of protein required for 50% inhibition of pyrogallol autoxidation per minute, the absorbance was measured at 440 nm in a microplate reader (Bio-Rad iMark™). To determine the enzymatic strength of catalase, 0.01M phosphate buffer solution (pH-7.0) and 0.2M H_2_O_2_ were added to the testicular protein homogenates. After thorough mixing, 5% potassium dichromate acetic acid reagent was added and the samples were kept in a boiling water bath for 10 minutes. As the blue colour mixtures turn into a green-coloured products of chromate acetate, the absorbance was measured at 570 nm. To measure the activities of the reduced glutathione (GSH), the protein samples were precipitated with 25% trichloroacetic acid (TCA) and spun down at 3000 rpm for 10 minutes at 4°C. The supernatant was mixed with 60 μM (5,5′-dithiobis-(2-nitrobenzoic acid) DTNB and 50 mM potassium phosphate buffer (pH-7.4). The absorbance of the resulting yellow-coloured reaction mixture was measured at 412 nm. To assay the activity of glutathione peroxidase (GPx), 100 μl of protein samples of testes were mixed with a neutral solution containing 0.32 M phosphate, 4 mM GSH, 10 mM sodium azide, 2.5 mM H_2_O_2_, 0.8mM EDTA. Then the tubes were incubated for 5 minutes at 37°C and centrifuged at 3500 rpm for 15 minutes. To the supernatant, 0.32M phosphate solution, and 60 μM DTNB reagent were added. GSH solutions corresponding to a concentration ranging between 4-20 μg/ml were also prepared and served as known standard. The intensity of yellow colour developed was measured in a microplate reader at 412 nm. Values were expressed as mg of GSH consumed per mg of protein [11].

### Statistical analysis

The values are represented as mean ± standard deviation. Student t test was applied to measure the statistical significance using Graph Pad Prism. The significance level was assumed at P < 0.05, unless otherwise indicated.

## Results

### BOTOX^®^ treatment protects against ageing mediated decline in total sperm count, motility and sperm morphological defects in experimental mice

The estimation of sperm count revealed a significant increase in the total number of sperms in the BOTOX^®^ treated group compared to the control group (Control: 9725000±2421260 vs BOTOX^®^: 15037500±2309897). The total percentage of the motile sperms (Control: 31.5±5.2 vs BOTOX^®^: 43.8±7.8) and the percentage of sperms with progressive motility (Control: 21.5±5.5 vs BOTOX^®^: 37.3±6.8) were found to be more in the BOTOX^®^ treated group when compared to that of control group (Fig 1). In the morphological analysis, the number of tailless sperms (Control: 16.5±4.8 vs BOTOX^®^: 6±1.5) as well as headless sperms (Control: 13.5±6.8 vs BOTOX^®^: 5±2.8) were found to be low in the BOTOX^®^ treated group when compared to control (Fig 2). However, no other obvious morphological differences of sperms were noticed between control and treatment group

**Fig 1.**
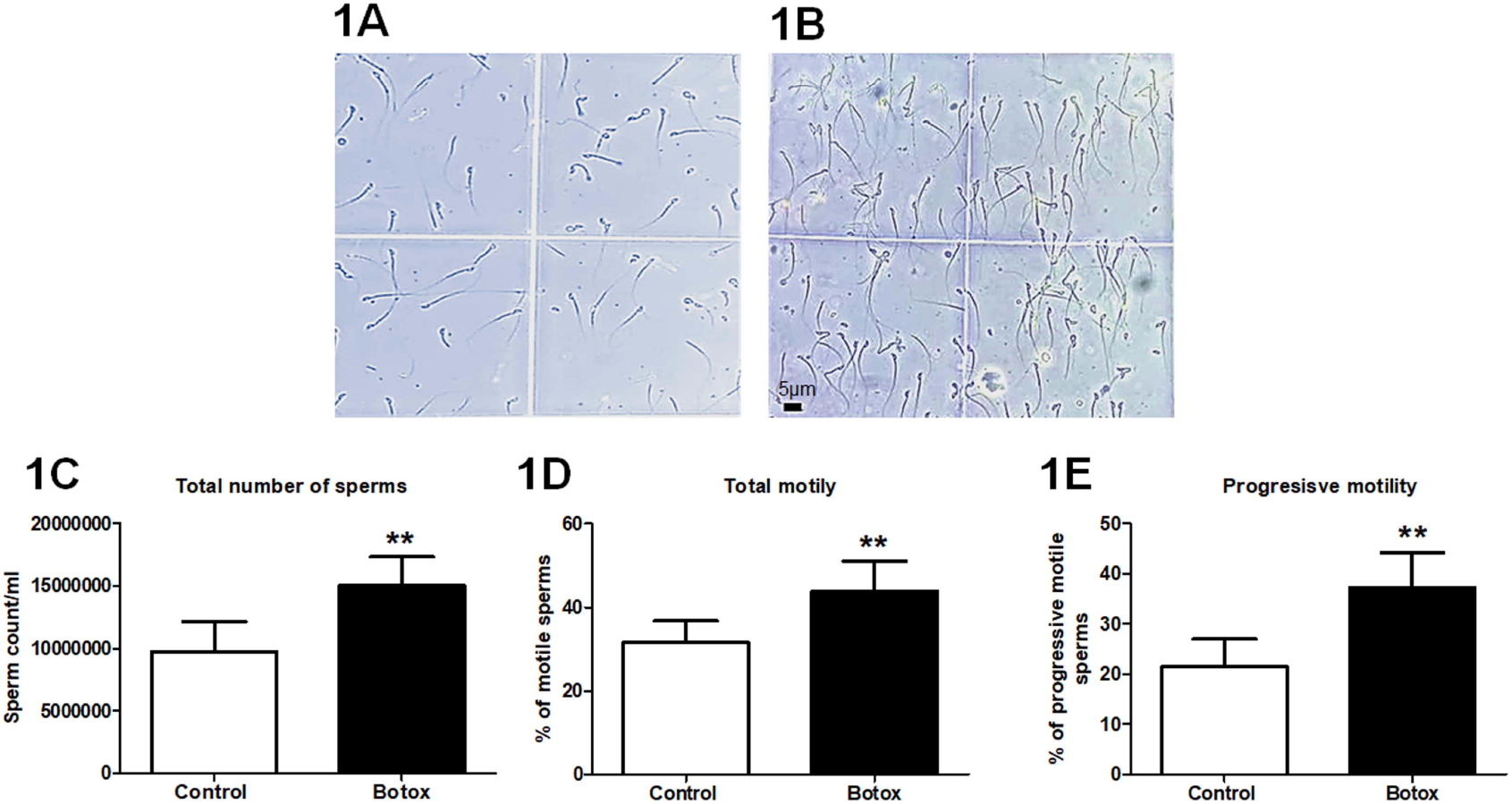
BOTOX^®^ treatment improves sperm count and motility. Phase contrast microscopy images of sperm isolated from the cauda epididymis from the control (1A) and BOTOX^®^ (1B) treated animals. The bar graphs represent the total sperm count (1C) and motility (1D) and progressive motility (1E) from the control and BOTOX^®^ treated animals.

**Fig 2.**
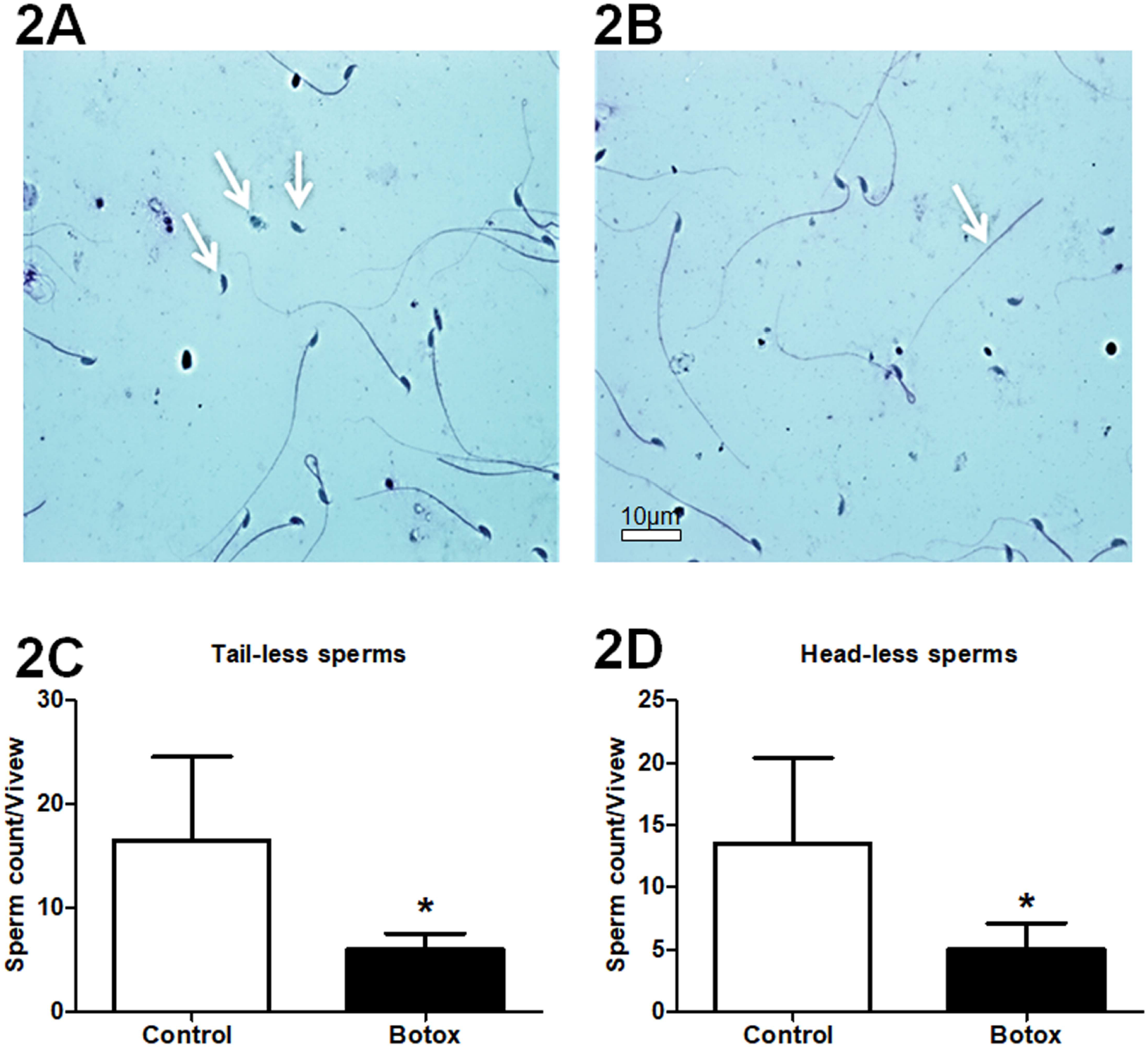
BOTOX^®^ treatment reduces the morphological defects of the sperm. The light microscopy images of tailless (2A) and headless (2B) sperm from control animals. The bar graphs represent the number of tailless sperms (2C) and headless (2D) sperms in the control and BOTOX^®^ treated animals.

### BOTOX^®^ treatment preserves the histological and spermatogenic capacity in ageing experimental mice

The histological examination of cross sections of the testes revealed no significant difference in the diameter of the seminiferous tubules in the testes between BOTOX® treated group and control group (Control: 21.8±6 vs BOTOX^®^: 22.6±2). Also, the diameter of the lumen of the seminiferous tubule was almost similar in the control and BOTOX^®^ treated animals (Control: 8.5±4.8 vs BOTOX^®^: 6.8±1.9) (Fig 3). The histological estimates of different cellular entities in the seminiferous tubules revealed a significant increase in the number of primary spermatocytes (Control: 39±3 vs BOTOX^®^: 49±6), secondary spermatocytes (Control: 20±3 vs BOTOX^®^: 28±2) and elongated spermatids (Control: 50±14 vs BOTOX^®^: 99±22) in the testis of the BOTOX^®^ treated group than that of the control group. However, the estimated number of round spermatids (Control: 86±17 vs BOTOX^®^: 63±16) was noticed to be reduced in the testis in the testis of the BOTOX^®^ treated group than control group (Fig 4).

**Fig 3.**
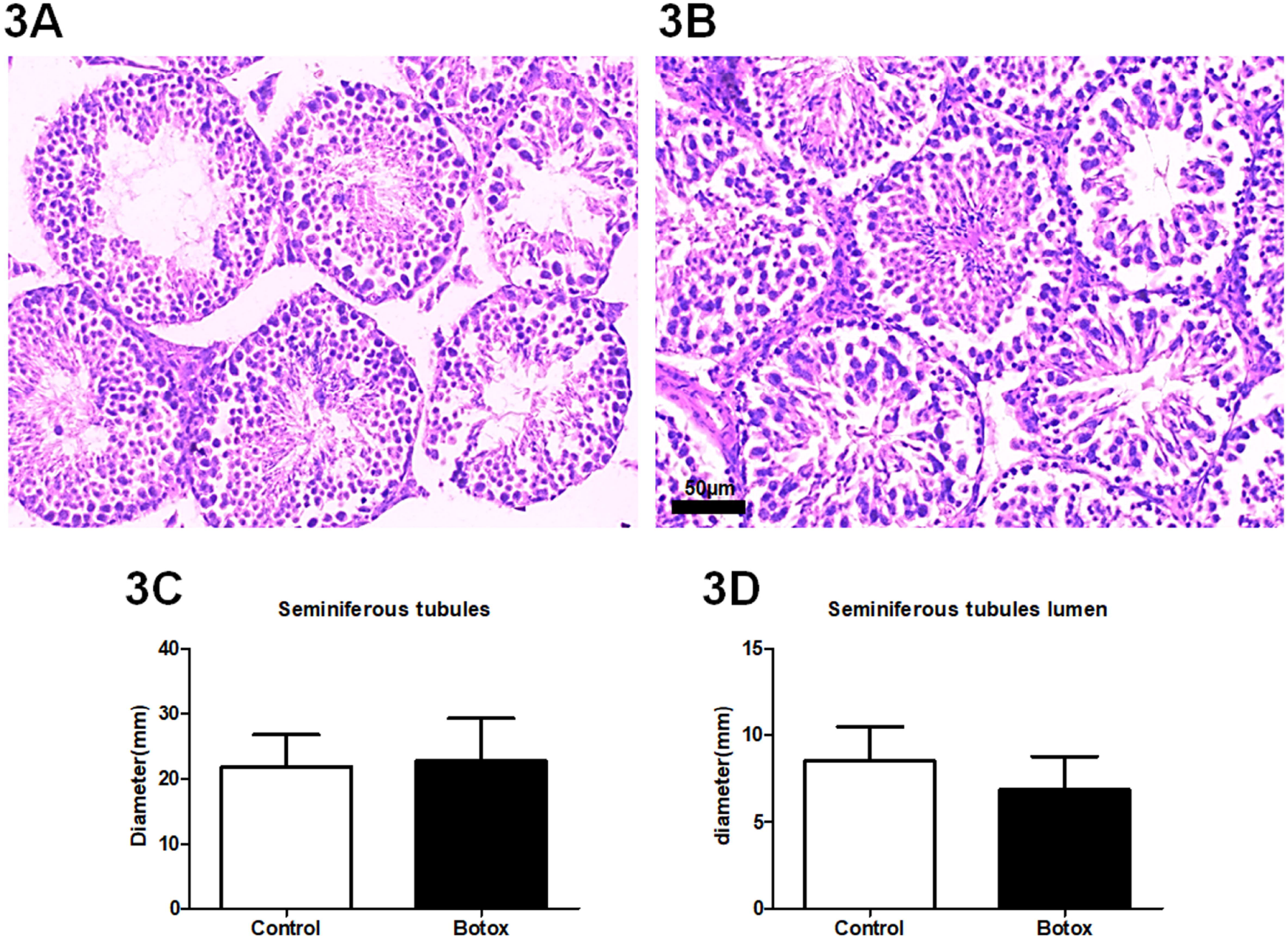
No major change in the structural difference in the seminiferous tubules between control and BOTOX^®^ treated animals. The H and E staining represent the morphology of seminiferous tubules in the testis of control (3A) and BOTOX^®^ treated animals (3B). The bar graph data represent the diameter of seminiferous tubules (3C) the diameter of lumen of seminiferous tubules (3D).

**Fig 4.**
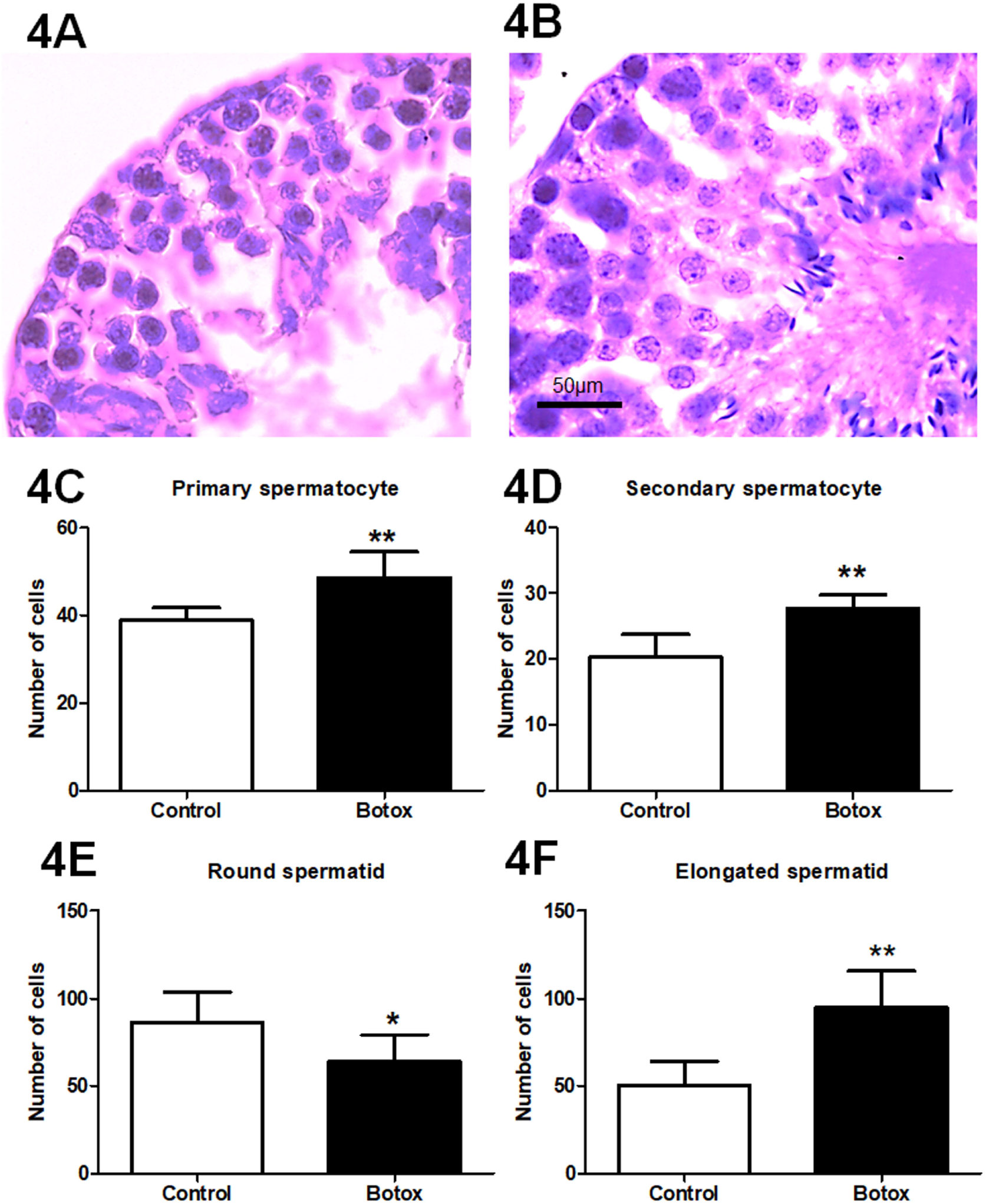
BOTOX^®^ treatment indicates increase of spermatogenesis. Light microscopic images of the H and E staining represent the cross sections of the seminiferous tubules in the control (4A) and BOTOX^®^ treated animals (4B). The bar graphs show the total numbers of primary spermatocytes (4D), secondary spermatocytes (4E), round spermatids (4F), and elongated spermatids (4G) per the cross section of the somniferous tubule in the testis of control and BOTOX® treated animals.

### BOTOX^®^ treated ageing experimental animals exhibit increased activities of testicular antioxidants

In the biochemical assessment of testicular protein extracts, the antioxidant activities of SOD (Control: 0.67±0.04 vs BOTOX^®^: 0.76±0.05), catalase (Control: 0.17±0.02 vs BOTOX^®^: 0.27±0.05), reduced glutathione (Control: 2.6. ±0.40 vs BOTOX^®^: 3.6±0.29), and glutathione peroxidase (Control: 0.40±0.004 vs BOTOX^®^ : 0.60±0.01) were considerably increased in the BOTOX^®^ treated group than that of the control group (Fig 5).

**Fig 5.**
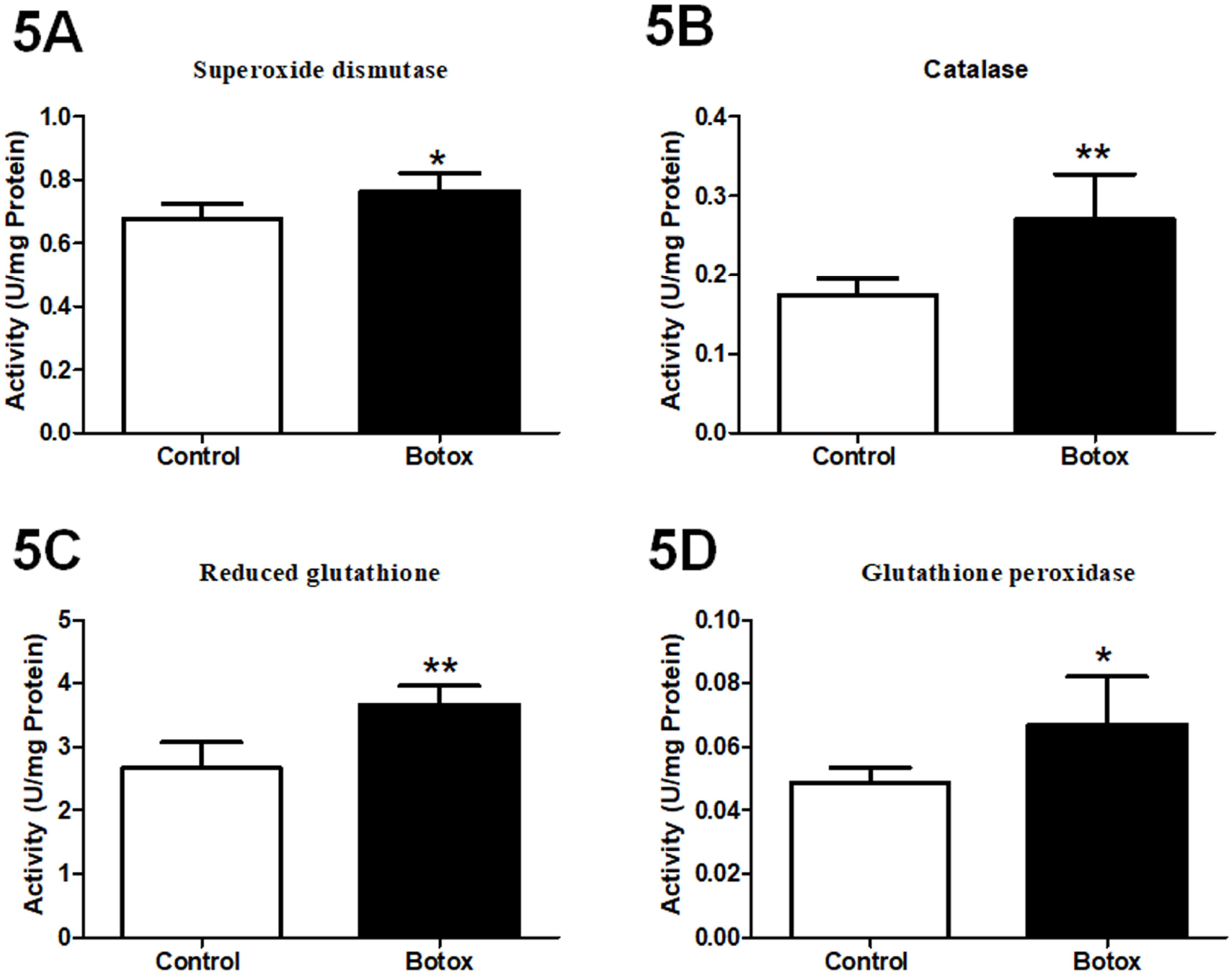
BOTOX^®^ treatment increases the activities of key antioxidants in the testis. The bar graph represents the antioxidant activity of SOD (5A), catalase (5B), reduced glutathione (5C) and glutathione peroxidase (5D) in U/mg protein from the testicular extracts of control and BOTOX^®^ treated animals.

## Discussion

The present study demonstrates that a trace amount of BOTOX^®^ injection at the distal intramuscular site in experimental ageing animals significantly improved the morphological and cytological characteristic of the seminiferous tubule in the testis and, quantity and motility of sperms in association with enhanced activities of testicular antioxidant enzymes. Recently, BOTOX^®^ has been reported to facilitate increased locomotion, cognitive functions and reduced anxiety related behaviours in ageing experimental rodents and humans [11,23,28,29]. It has been reported that BOTOX^®^ contributes to the improved blood circulation and oxygen supply leading to trophic support to cell cycle events and cryoprotection [30,31]. Considering the fact, intramuscular injection of BOTOX^®^ can be predicted to enhance the blood circulation in the testis of the experimental animals which can be expected to promote the histological parameters and increased number of key cellular compartments of the testis. While the BOTOX^®^ treated animals showed a significant increase in the number of primary and secondary spermatocytes, and elongated spermatids, a reduction in the number of round spermatids was evident in BOTOX^®^ treated animals. The latter could be due to the accelerated differentiation of round to elongated spermatid contributing to an enhanced spermiogenesis in the BOTOX^®^ treated group, as the therapeutic botulinum toxin has been reported to facilitate cellular differentiation of different tissues [32,33].

Spermiogenesis is the cellular process of the testis by which haploid round spermatids undergo a series of events to become motile spermatozoa [34,35]. Spermiogenesis commences after spermatocytes have accomplished the meiotic reductive cycle [36]. This process involves multifaceted morphological, biochemical, and physiological alterations in spermatids [34,36,37]. The major events in this process involve embellishment of the acrosome from the Golgi apparatus, condensation of the chromatin, centrosome disintegration, elimination of redundant cytoplasmic portion and formation of flagellum responsible for the motility of the spermatozoa [38,39]. Thus, the molecular changes associated with BOTOX® treatment may involve the progression of the flagellum, differentiation of spermatozoa and increased motility of sperm in the testis.

While physiological levels of ACh are important for spermatogenesis and motility of sperms [40], abnormal levels of ACh appear to induce oxidative stress in the testis and reduce the motility and function of sperms [41,42]. Oxidative stress has mostly been associated with loss of structural and functional integrity of spermatozoa by inducing damage to DNA, RNA transcripts and telomeres thereby leading to testicular atrophy and infertility in ageing and disease [43,44]. Oxidative stress in the testis contributes to low sperm count, motility and abnormal morphology [45,46]. Earlier studies have indicated that the supplementation of antioxidants like vitamin C, vitamin E and glutathione improve spermatogenesis and sperm quality [47,48]. Recent studies have demonstrated that the use of therapeutic botulinum toxin reduced oxidative stress in animal models and plasma of patients [22,49,50], Yesudhas et al. has recently reported that BOTOX® improves the antioxidant enzyme activity and provide neuroprotection in the brain of ageing experimental animals [11,23]. Considering the fact, BOTOX^®^ treatment could potentially elicit cytotropic signalling events or neutralize the apoptotic signalling cascades in the testis leading to the increased survival of cellular components of the seminiferous tubules.

Favaretto et,al. reported that abnormal levels of endogenous ACh can interfere with the production of testosterone secretion in rat Leydig cells [51]. Similarly, cholinergic agonists have been reported to mediate inhibition testosterone biosynthesis in the testis [52]. Therefore, BOTOX^®^ might counteract the ageing associated excessive release of ACh and regulate the production of testosterone and antioxidant enzymes thereby facilitating spermatogenesis. Taken together, a mild amount of BOTOX^®^ might be a potential therapeutic aid to treat male infertility.

## Conclusion

Reduced sperm count responsible for male infertility has been a rising medical concern worldwide. The risk for male infertility includes ageing, familial, sporadic and abnormal lifestyle, environmental factors, testicular damage and atrophy, infection, exposure to radiation, chemicals, pesticides, hazardous materials, occupational hazards, alcohol consumption, drug addiction, disorders with metabolic, vascular, neurological, psychiatric and malignancy complications. Though hormonal replacement, aromatase inhibitors and dopamine antagonists, and some surgical procedures have been practised to correct male infertility, the efficacy of these treatments has been a subject of debate and often these treatments appear to pose adverse effects than clinical rectification. Recently, cholinergic toxicity has been proposed as an underlying cause of erectile dysfunction, testicular atrophy and infertility. Recently, blockade of ACh release by BOTOX^®^ has been used to treat various testicular defects like bilateral cremasteric muscle spasms, retractile testis, cryptorchidism and erectile dysfunction. The present study demonstrates that a mild intramuscular injection of BOTOX^®^ provides defends against ageing mediated spermatogenic decline and improves the total sperm count and motility in correlation with increased activities of key antioxidants in the testis of experiment mice. Therefore, BOTOX^®^ might be considered as a potential drug to treat male infertility. However, possibilities for the unknown adverse effects associated with BOTOX^®^ might not completely be excluded.

## Declaration of Competing Interest

The authors declare that they have no conflict of interest.

## Acknowledgement

MK has been supported by the Faculty Recharge Programme, University Grants Commission (UGC-FRP), New Delhi, India. This work has been supported by a research grant (SERB-EEQ/2016/000639) and an Early Career Research Award (SERB-ECR/2016/000741) from Science and Engineering Research Board (SERB) under the Department of Science and Technology (DST), Government of India. RKR was supported as JRF from the project grant-ECR/2016/000741, SERB, India. The authors acknowledge, RUSA2.0, Biological Sciences, BDU for the financial support, UGC-SAP, DST-FIST for the infrastructure of the Department of Animal Science, Bharathidasan University. The authors would like to thank Dr. Muthuswamy Anusuyadevi for helpful advice and support on biochemical assays.

## Authors’ contribution

MK conceived the idea, design of the study. RKR and BK contributed to further development of the concept and design of the study RKR, AS SR, and MK were involved in the acquisition of experimental data and analysis. MK and RKR were involved in the initial draft of the manuscript. All authors contributed to the further revision of article, made critical comments and suggestions. All authors discussed the content and approved the final version manuscript.

## Notes

### Competing Interest Statement

The authors have declared no competing interest.

## Reference

[1] Picciotto MR, Higley MJ, Mineur YS. Acetylcholine as a neuromodulator: cholinergic signaling shapes nervous system function and behavior. Neuron 2012;76:116–29. https://doi.org/10.1016/j.neuron.2012.08.036.

[2] Han X, Zhang C, Ma X, Yan X, Xiong B, Shen W, et al. Muscarinic acetylcholine receptor M5 is involved in spermatogenesis through the modification of cell–cell junctions. Reproduction 2021;162:47–59. https://doi.org/10.1530/REP-21-0079.

[3] Schirmer SU, Eckhardt I, Lau H, Klein J, DeGraaf YC, Lips KS, et al. The cholinergic system in rat testis is of non-neuronal origin. Reproduction 2011;142:157–66. https://doi.org/10.1530/REP-10-0302.

[4] Arıcan EY, Gökçeoğlu Kayalı D, Ulus Karaca B, Boran T, Öztürk N, Okyar A, et al. Reproductive effects of subchronic exposure to acetamiprid in male rats. Sci Rep 2020;10:8985. https://doi.org/10.1038/s41598-020-65887-0.

[5] da Silva Júnior ED, de Souza BP, Rodrigues JQD, Caricati-Neto A, Jurkiewicz A, Jurkiewicz NH. Functional characterization of acetylcholine receptors and calcium signaling in rat testicular capsule contraction. Eur J Pharmacol 2013;714:405–13. https://doi.org/10.1016/j.ejphar.2013.07.007.

[6] Tata AM, Velluto L, D’Angelo C, Reale M. Cholinergic system dysfunction and neurodegenerative diseases: cause or effect? CNS Neurol Disord Drug Targets 2014;13:1294–303. https://doi.org/10.2174/1871527313666140917121132.

[7] Zhu C, Palmada MN, Aguado LI, Cavicchia JC. Administration of acetylcholine to the spermatic nerve plexus inhibits testosterone secretion in an in vitro isolated rat testis-nerve plexus system. Int J Androl 2002;25:134–8. https://doi.org/10.1046/j.1365-2605.2002.00337.x.

[8] Kandasamy M. Perspectives for the use of therapeutic Botulinum toxin as a multifaceted candidate drug to attenuate COVID-19. Med Drug Discov 2020;6:100042. https://doi.org/10.1016/j.medidd.2020.100042.

[9] Nigam PK, Nigam A. BOTULINUM TOXIN. Indian J Dermatol 2010;55:8–14. https://doi.org/10.4103/0019-5154.60343.

[10] Pirazzini M, Rossetto O, Eleopra R, Montecucco C. Botulinum Neurotoxins: Biology, Pharmacology, and Toxicology. Pharmacol Rev 2017;69:200–35. https://doi.org/10.1124/pr.116.012658.

[11] Yesudhas A, Radhakrishnan RK, Sukesh A, Ravichandran S, Manickam N, Kandasamy M. BOTOX® counteracts the innate anxiety-related behaviours in correlation with increased activities of key antioxidant enzymes in the hippocampus of ageing experimental mice. Biochem Biophys Res Commun 2021;569:54–60. https://doi.org/10.1016/j.bbrc.2021.06.071.

[12] Breikaa RM, Mosli HA, Abdel-Naim AB. Influence of Onabotulinumtoxin A on testes of the growing rat. J Biochem Mol Toxicol 2016;30:608–13. https://doi.org/10.1002/jbt.21828.

[13] Breikaa RM, Mosli HA, Nagy AA, Abdel-Naim AB. Adverse testicular effects of Botox® in mature rats. Toxicol Appl Pharmacol 2014;275:182–8. https://doi.org/10.1016/j.taap.2014.01.003.

[14] Ramelli E, Brault N, Tierny C, Atlan M, Cristofari S. Intrascrotal injection of botulinum toxin A, a male genital aesthetic demand: Technique and limits. Progrès En Urologie 2020;30:312–7. https://doi.org/10.1016/j.purol.2020.04.016.

[15] Cousins E, Ward A, Roffe C, Rimington L, Pandyan A. Does low-dose botulinum toxin help the recovery of arm function when given early after stroke? A phase II randomized controlled pilot study to estimate effect size. Clin Rehabil 2010;24:501–13. https://doi.org/10.1177/0269215509358945.

[16] Padda IS, Tadi P. Botulinum Toxin. StatPearls, Treasure Island (FL): StatPearls Publishing; 2021.

[17] Wissel J, Heinen F, Schenkel A, Doll B, Ebersbach G, Müller J, et al. Botulinum toxin A in the management of spastic gait disorders in children and young adults with cerebral palsy: a randomized, double-blind study of “high-dose” versus “low-dose” treatment. Neuropediatrics 1999;30:120–4. https://doi.org/10.1055/s-2007-973475.

[18] Ghanem H, Raheem AA, AbdelRahman IFS, Johnson M, Abdel-Raheem T. Botulinum Neurotoxin and Its Potential Role in the Treatment of Erectile Dysfunction. Sex Med Rev 2018;6:135–42. https://doi.org/10.1016/j.sxmr.2017.07.008.

[19] Raef HS, Elmariah SB. Treatment of male genital dysesthesia with botulinum toxin. JAAD Case Rep 2021;10:60–2. https://doi.org/10.1016/j.jdcr.2021.01.018.

[20] Reddy AG, Dick BP, Natale C, Akula KP, Yousif A, Hellstrom WJG. Application of Botulinum Neurotoxin in Male Sexual Dysfunction: Where Are We Now? Sex Med Rev 2021;9:320–30. https://doi.org/10.1016/j.sxmr.2020.05.004.

[21] Sekiguchi A, Motegi S-I, Uchiyama A, Uehara A, Fujiwara C, Yamazaki S, et al. Botulinum toxin B suppresses the pressure ulcer formation in cutaneous ischemia-reperfusion injury mouse model: Possible regulation of oxidative and endoplasmic reticulum stress. J Dermatol Sci 2018;90:144–53. https://doi.org/10.1016/j.jdermsci.2018.01.006.

[22] Uchiyama A, Yamada K, Perera B, Ogino S, Yokoyama Y, Takeuchi Y, et al. Protective effect of botulinum toxin A after cutaneous ischemia-reperfusion injury. Sci Rep 2015;5:9072. https://doi.org/10.1038/srep09072.

[23] Yesudhas A, Roshan SA, Radhakrishnan RK, Abirami GPP, Manickam N, Selvaraj K, et al. Intramuscular Injection of BOTOX® Boosts Learning and Memory in Adult Mice in Association with Enriched Circulation of Platelets and Enhanced Density of Pyramidal Neurons in the Hippocampus. Neurochem Res 2020;45:2856–67. https://doi.org/10.1007/s11064-020-03133-9.

[24] Kanimozhi V, Palanivel K, Kadalmani B, Krikun G, Taylor HS. Apolipoprotein E Induction in Syrian Hamster Testis Following Tributyltin Exposure: A Potential Mechanism of Male Infertility. Reprod Sci 2014;21:1006–14. https://doi.org/10.1177/1933719114522519.

[25] Mehraein F, Negahdar F. Morphometric evaluation of seminiferous tubules in aged mice testes after melatonin administration. Cell J 2011;13:1–4.

[26] Nakata H, Wakayama T, Takai Y, Iseki S. Quantitative Analysis of the Cellular Composition in Seminiferous Tubules in Normal and Genetically Modified Infertile Mice. J Histochem Cytochem 2015;63:99–113. https://doi.org/10.1369/0022155414562045.

[27] Lowry OH, Rosebrough NJ, Farr AL, Randall RJ. Protein measurement with the Folin phenol reagent. J Biol Chem 1951;193:265–75.

[28] de Jongh R, Bolt I, Schermer M, Olivier B. Botox for the brain: enhancement of cognition, mood and pro-social behavior and blunting of unwanted memories. Neurosci Biobehav Rev 2008;32:760–76. https://doi.org/10.1016/j.neubiorev.2007.12.001.

[29] Lewis MB, Bowler PJ. Botulinum toxin cosmetic therapy correlates with a more positive mood. J Cosmet Dermatol 2009;8:24–6. https://doi.org/10.1111/j.1473-2165.2009.00419.x.

[30] Schweizer DF, Schweizer R, Zhang S, Kamat P, Contaldo C, Rieben R, et al. Botulinum toxin A and B raise blood flow and increase survival of critically ischemic skin flaps. J Surg Res 2013;184:1205–13. https://doi.org/10.1016/j.jss.2013.04.004.

[31] Zhibo X, Miaobo Z. Botulinum toxin type A affects cell cycle distribution of fibroblasts derived from hypertrophic scar. J Plast Reconstr Aesthet Surg 2008;61:1128–9. https://doi.org/10.1016/j.bjps.2008.05.003.

[32] Gugerell A, Kober J, Schmid M, Nickl S, Kamolz LP, Keck M. Botulinum toxin A and lidocaine have an impact on adipose-derived stem cells, fibroblasts, and mature adipocytes in vitro. J Plast Reconstr Aesthet Surg 2014;67:1276–81. https://doi.org/10.1016/j.bjps.2014.05.029.

[33] Jeong HS, Lee BH, Sung HM, Park SY, Ahn DK, Jung MS, et al. Effect of Botulinum Toxin Type A on Differentiation of Fibroblasts Derived from Scar Tissue. Plast Reconstr Surg 2015;136:171e–8e. https://doi.org/10.1097/PRS.0000000000001438.

[34] Linn E, Ghanem L, Bhakta H, Greer C, Avella M. Genes Regulating Spermatogenesis and Sperm Function Associated With Rare Disorders. Front Cell Dev Biol 2021;9:634536. https://doi.org/10.3389/fcell.2021.634536.

[35] O’Donnell L. Mechanisms of spermiogenesis and spermiation and how they are disturbed. Spermatogenesis 2015;4:e979623. https://doi.org/10.4161/21565562.2014.979623.

[36] Griswold MD. Spermatogenesis: The Commitment to Meiosis. Physiol Rev 2016;96:1–17. https://doi.org/10.1152/physrev.00013.2015.

[37] Dadoune JP. The cellular biology of mammalian spermatids: a review. Bull Assoc Anat (Nancy) 1994;78:33–40.

[38] Berruti G, Paiardi C. Acrosome biogenesis: Revisiting old questions to yield new insights. Spermatogenesis 2011;1:95–8. https://doi.org/10.4161/spmg.1.2.16820.

[39] Suphamungmee W, Wanichanon C, Vanichviriyakit R, Sobhon P. Spermiogenesis and chromatin condensation in the common tree shrew, Tupaia glis. Cell Tissue Res 2008;331:687–99. https://doi.org/10.1007/s00441-007-0557-5.

[40] Bray C, Son J-H, Meizel S. Acetylcholine causes an increase of intracellular calcium in human sperm. Mol Hum Reprod 2005;11:881–9. https://doi.org/10.1093/molehr/gah245.

[41] Ngoula F, Watcho P, Dongmo M-C, Kenfack A, Kamtchouing P, Tchoumboué J. Effects of pirimiphos-methyl (an organophosphate insecticide) on the fertility of adult male rats. Afr Health Sci 2007;7:3–9.

[42] Slimen S, Saloua EF, Najoua G. Oxidative stress and cytotoxic potential of anticholinesterase insecticide, malathion in reproductive toxicology of male adolescent mice after acute exposure. Iran J Basic Med Sci 2014;17:522–30.

[43] Aitken RJ, Gibb Z, Baker MA, Drevet J, Gharagozloo P. Causes and consequences of oxidative stress in spermatozoa. Reprod Fertil Dev 2016;28:1–10. https://doi.org/10.1071/RD15325.

[44] Sabeti P, Pourmasumi S, Rahiminia T, Akyash F, Talebi AR. Etiologies of sperm oxidative stress. Int J Reprod Biomed 2016;14:231–40.

[45] Agarwal A, Virk G, Ong C, du Plessis SS. Effect of Oxidative Stress on Male Reproduction. World J Mens Health 2014;32:1–17. https://doi.org/10.5534/wjmh.2014.32.1.1.

[46] Alahmar AT. Role of Oxidative Stress in Male Infertility: An Updated Review. J Hum Reprod Sci 2019;12:4–18. https://doi.org/10.4103/jhrs.JHRS_150_18.

[47] Ahmadi S, Bashiri R, Ghadiri-Anari A, Nadjarzadeh A. Antioxidant supplements and semen parameters: An evidence based review. Int J Reprod Biomed 2016;14:729–36.

[48] Majzoub A, Agarwal A. Systematic review of antioxidant types and doses in male infertility: Benefits on semen parameters, advanced sperm function, assisted reproduction and live-birth rate. Arab J Urol 2018;16:113–24. https://doi.org/10.1016/j.aju.2017.11.013.

[49] Dini E, Mazzucchi S, De Luca C, Cafalli M, Chico L, Lo Gerfo A, et al. Plasma Levels of Oxidative Stress Markers, before and after BoNT/A Treatment, in Chronic Migraine. Toxins (Basel) 2019;11:E608. https://doi.org/10.3390/toxins11100608.

[50] Zhou Y, Yu S, Zhao J, Feng X, Zhang M, Zhao Z. Effectiveness and Safety of Botulinum Toxin Type A in the Treatment of Androgenetic Alopecia. Biomed Res Int 2020;2020:1501893. https://doi.org/10.1155/2020/1501893.

[51] Favaretto AL, Valença MM, Picanço-Diniz DL, Antunes-Rodrigues JA. Inhibitory role of cholinergic agonists on testosterone secretion by purified rat Leydig cells. Arch Int Physiol Biochim Biophys 1993;101:333–5. https://doi.org/10.3109/13813459309046988.

[52] Kasson BG, Hsueh AJ. Nicotinic cholinergic agonists inhibit androgen biosynthesis by cultured rat testicular cells. Endocrinology 1985;117:1874–80. https://doi.org/10.1210/endo-117-5-1874.

